# Apicomplexan motility depends on the operation of an endocytic-secretory cycle

**DOI:** 10.1101/450080

**Authors:** Simon Gras, Elena Jimenez-Ruiz, Christen M. Klinger, Leandro Lemgruber, Markus Meissner

## Abstract

Apicomplexan parasites invade host cells in an active process, involving their ability to move by gliding motility and invasion. While the acto-myosin-system of the parasite plays a crucial role in the formation and release of attachment sites during this process, there are still open questions, such as how the force powering motility is generated. In many eukaryotes a secretory-endocytic cycle leads to recycling of receptors (integrins), necessary to form attachment sites, regulation of surface area during motility and generation of retrograde membrane flow. Here we demonstrate that endocytosis operates during gliding motility in *Toxoplasma gondii* and appears to be crucial for the establishment of retrograde membrane flow, since inhibition of endocytosis blocks retrograde flow and motility. We identified lysophosphatidic acid (LPA) as a potent stimulator of endocytosis and demonstrate that extracellular parasites can efficiently incorporate exogenous material, such as nanogold particles. Furthermore, we show that surface proteins of the parasite are recycled during this process. Interestingly, the endocytic and secretory pathways of the parasite converge, and endocytosed material is subsequently secreted, demonstrating the operation of an endocytic-secretory cycle. Together our data consolidate previous findings and we propose a novel model that reconciles parasite motility with observations in other eukaryotes: the fountain-flow-model for apicomplexan parasite motility.

## INTRODUCTION

The intracellular protozoan parasite *Toxoplasma gondii* infects nearly 2 billion people globally. This apicomplexan can cause severe disease in immunocompromised people and can lead to miscarriage or malformation of the foetus in pregnant women (1). During the acute phase of infection, the tachyzoite rapidly replicates inside the host cell within a specialised compartment, the parasitophorous vacuole, which itself is formed during active invasion (2).

Like all apicomplexans, *T. gondii* invades host cells in an active process, involving both the parasite’s ability to move by gliding motility and invasion factors derived from the unique secretory organelles localised at the parasite’s apical pole (micronemes and rhoptries) (3). According to the linear motor model, micronemal transmembrane proteins are secreted at the apical tip of the parasite and act as force transmitters by interacting with both the substrate and the acto-myosin system of the parasite. However, recent studies suggested a different role of the acto-myosin system during motility, since parasites devoid of F-actin are significantly reduced in overall motility, but still capable of moving at a similar speed as wild-type parasites. This surprising finding has been attributed to a function of the acto-myosin as a surface clutch that allows force transmission by regulating the formation and release of attachment sites, akin to other eukaryotes (4). In good agreement with this hypothesis, mutants for this system cannot efficiently attach to the substrate (5-8).

During motility, most eukaryotic cells show a capping activity of surface ligands, which is dependent on actin, microtubules, and a secretory-endocytic cycle, leading to the establishment of a retrograde membrane flow (4, 9). A recent study on *Dictyostelium* provided direct evidences for the fluid flow model during cell migration (10). This study demonstrated that, during migration of *Dictyostelium*, the membrane volume of the cell remains constant due to the occurrence of a secretory-endocytic cycle. This circulation follows a fountain flow model, where new membrane lipids are delivered to the anterior cell membrane, whereas excess membrane is recycled at the rear. Interestingly, in this study a direct relationship between cell migration and membrane turnover rate was observed, suggesting that the cells establish a fluid drive that contributes to the generation of force required for motility, as suggested previously (4). Importantly, it appears that myosin and actin have only a supporting function in the establishment of the fluid drive, since treatment with actin-or myosin-disrupting drugs, such as latrunculin B or blebbistatin, did not significantly affect membrane movement (10). In good agreement, retrograde membrane flow in apicomplexan parasites is not strictly dependent on parasite actin, as shown for *Plasmodium* sporozoites and *Toxoplasma* tachyzoites (7, 11).

It is well accepted that gliding motility of apicomplexans depends on the regulated secretion of the apically localised micronemes (12). While this dependency was previously contributed to the secretion of surface ligands, such as MIC2, that are required as force transmitters, it is also possible that polarised secretion is required for the generation of retrograde membrane flow, akin to the fountain flow model (10) and as previously suggested for *T. gondii* (7, 13). However, to date it is not fully understood how apicomplexan parasites ensure a constant cell surface during motility by removing excess membrane deposited on the surface due to microneme secretion.

While the previously described shedding of membrane trails during motility (14) might contribute to a constant membrane content and cell surface, it appears likely that, as suggested by the fountain flow model (10), excess membrane is internalised and recycled during motility. Besides, if shedding was the only mechanism to achieve membrane balance, the parasite would be forced to synthesize an energetically unfeasible quantity of new membrane lipids/proteins, which seems unrealistic for a parasite with limited metabolic abilities.

Here we set out to determine if extracellular parasites are capable of efficiently recycling membrane and taking up exogenous material via endocytosis. To date, uptake of exogenous material has been demonstrated during the intracellular stages of the parasite (15, 16). In other eukaryotes, endocytic processes play key roles in membrane dynamics, making it an important participant in cell motility (17, 18). Endocytosis can occur via different mechanisms and are roughly defined as CDE (clathrin-dependent endocytosis) or CIE (clathrin-independent endocytosis) (19). Apicomplexan genomes lack many factors known to be involved in the endocytic system, such as the ESCRT complex, and previous reverse genetic analysis suggested that the remaining factors were repurposed to contribute to the biogenesis and maintenance of unique organelles, such as the Inner Membrane Complex (IMC) or the secretory organelles (20-25).

Here, we demonstrate the implication of endocytosis in the maintenance of retrograde membrane flow and provide a link between this process and gliding motility, in good agreement with the fountain flow model (4, 10). We demonstrate the capacity of extracellular tachyzoites to take up phospholipids, 10 nm nanogold particles, and antibodies directed against parasite surface proteins. Interestingly, endocytic uptake of material appears to occur in a clathrin-independent manner and follows the known secretory pathway of the parasite, with accumulation of material in the rhoptries but also vacuolar-like compartment (VAC, PLV, (26, 27)).

Together our data demonstrate the existence of an secretory-endocytic cycle during parasite motility that appears capable of generating the force for motility and therefore fully supports the hypothesis of a fountain-flow model, as suggested for other motile eukaryotic cells (10).

## RESULTS

### Fountain flow model and evidence of endocytosis implication in *T. gondii* motility

The fountain flow model has been recently demonstrated to operate during eukaryotic cell motility, such as in *Dictyostelum discoideum* (10). This model predicts the establishment of a retrograde membrane flow by localised secretion (at the anterior end of the cell), followed by endocytic recycling to ensure membrane balance. In this respect, apicomplexan parasites are a prime example of highly polarised cells, where the micronemes are secreted at the apical tip during gliding (Fig.1A), which, in analogy to *D. discoideum*, should result in retrograde membrane flow.

**Figure 1:**
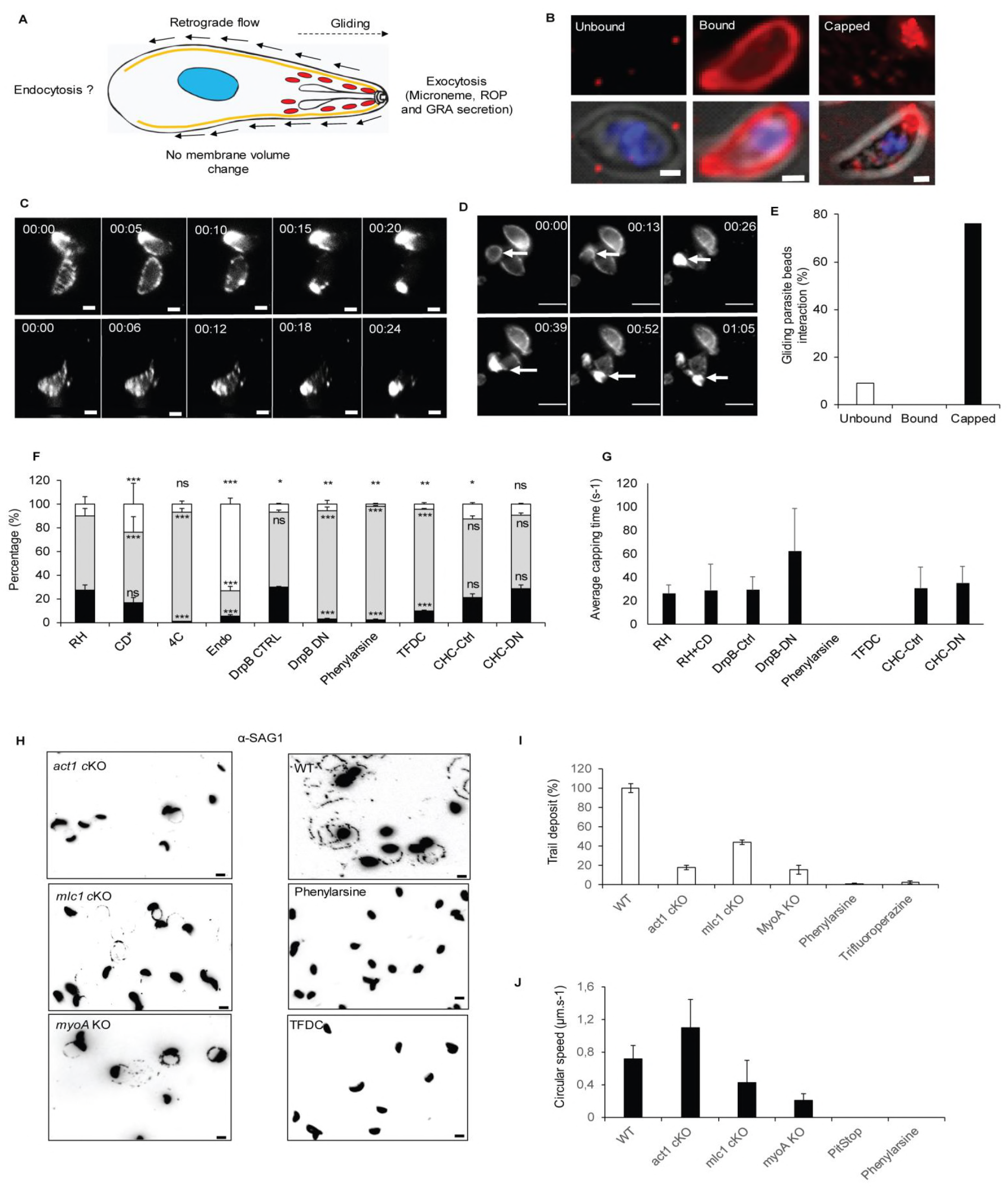
Evidence for involvement of endocytosis in *T. gondii* motility. (A): Fountain flow model described for *Dictyostelium* by Takana *et al.* 2017 applied to *T. gondii*. While it was demonstrated that motility depends on apical secretion of micronemes, the role of endocytosis to ensure membrane balance and recycling is unclear (B): Representative pictures of the three phenotypes observed in the bead translocation experiment: Unbound: parasite without beads, that either did not interact or lost their interaction with the beads; Bound: parasite with beads around the plasma membrane; Capped: parasite which translocated bound beads to its basal pole. Scale bar 1µm. (C): Time lapse analysis of bead translocation. Parasites were incubated with latex beads. Capping was recorded by live microscopy to determine the average time required for capping. Scale bar 1 µm. (D): Example of a parasite that translocates beads and initiates gliding motility. Scale bar 5µM. (E): Quantification of gliding parasites in combination with bead translocation. Only parasites with un bound (No beads) and capped beads (Beads accumulated at the posterior end) were observed gliding. No instances where beads were bound at the plasma membrane, but not capped, were observed. n = 38. (F): Quantification of bead translocation using indicated inhibitors or parasite mutants. Un-bound (*white*), bound (*grey*) and capped (*black*). Parasites expressing dominant negative versions of DrpB or CHC1 were induced with 1 µM Shield-1 as described previously (21, 23). Inhibitors were used at the following concentrations: 0.5 μM Cytochalasin-D (CD), 10µM phenyl arsine oxide or 50µM Trifluoperazine dihydrochloride (TFDC). Mean values of three independent assays are shown ± SEM. *** *p*-value <0.001 in a two-tailed Student’s t-test compared to RH without inhibitors. (G): Analysis of the average time required for capping under the conditions shown in (F). Analysis in the presence of endocytosis inhibitors was not possible, since no capping could be observed under these conditions. (H-I): Trail deposition assay of wild-type (WT) parasites compared to motor mutants (*act1* cKO, *mlc1* cKO, and *myoA* KO) and endocytosis inhibitors (phenylarsine and TFDC). (H), representative pictures, scale bar 5 µm. (I), Quantification of the trail deposition. Mean values of three independent assays are shown ± SEM. (J): Analysis of average gliding speed. Mean values of three independent assays are shown ± SEM.

To analyse retrograde membrane flow in *T. gondii* tachyzoites we previously adapted a translocation assay (28) that follows the transport of beads to the posterior end of the parasite. This ‘capping’ has been directly implicated in the motility of *T. gondii*, and, importantly, can occur in the absence of the acto-myosin motor (7). Three phenotypes are observable during this assay: “Unbound” parasites without beads, “Bound” parasites with beads distributed throughout the plasma membrane, and “Capped” parasites that have translocated all the bead to their basal pole (Fig. 1B).

We performed live imaging of the capping process in the presence and absence of different inhibitors or proteins that interfere with the acto-myosin system, secretion, or endocytic processes in other eukaryotes (Fig.1C, Video S1). We incubated parasites with 40 nm latex beads at 4°C to allow binding of the beads to the parasites surface. Upon a temperature shift to 37°C capping occurs rapidly, and beads accumulate at the posterior pole of the parasite within ∼25 seconds (Fig. 1C). Importantly, we found that parasite gliding correlates with bead translocation (Video S2), since ∼76% of motile parasites showed translocation, while ∼24% of motile parasites did not have any beads on the surface, indicating that no initial binding of the beads occurred or that beads were shed after translocation, as seen in movie S3. Importantly, no gliding parasites were identified where beads remained immobile while bound to the parasite plasma membrane (Fig.1B, D, E). Interestingly, parasites can translocate beads without moving, demonstrating that retrograde flow can occur in the absence of gliding, while gliding does not occur in the absence of retrograde flow.

The mechanism underlying parasite membrane balance is unknown and suggested to depend exclusively on membrane shedding and processing of micronemal transmembrane proteins (29). We hypothesised that, akin to other eukaryotes, membrane balance is also ensured by endocytic recycling of excess membrane and proteins (10). To determine if *T. gondii* retrograde flow could be dependent on a similar mechanism, we tested conditions that inhibit or alter secretion/exocytosis using established inhibitors of endocytosis or parasite strains, such as parasites expressing dominant negative (DN) DrpB (DrpB-DN) (23), where micronemes are absent (Figure 1F).

Parasites incubated on ice bound beads (92.5±2.6%), but no translocation was observed; a temperature shift to 37°C resulted in translocation in ∼27% of parasites. Interestingly, incubation of parasites in the presence of 0.5 µM Cytochalasin D (CD, a drug used to disrupt F-actin) did not result in significant reduction of bead translocation, confirming that retrograde membrane flow can occur in the absence of a functional acto-myosin system, as reported previously (7). In sharp contrast, abrogation of microneme secretion, either by incubation of parasites in endo buffer (2) or induction of the DrpB-DN strain, abolishes bead translocation, demonstrating that retrograde flow depends on polarised secretion as predicted by the fountain flow model (Fig.1A; (10). To evaluate if capping could depend on endocytosis, we used well established inhibitors of endocytosis, such as Phenylarsine oxide (30) and Trifluoroperazine (31), as well as a DN strain for clathrin heavy chain (CHC-Hub) (21). While the endocytosis inhibitors abrogated capping (2.3±0.33% and 9.8±0.83% capping respectively), expression of dominant negative CHC1 did not result in significant reduction of capping (-Shield: 21.3±3.1%, +Shield: 28.9±2.6% capping). Together, these results suggest that retrograde membrane flow depends on an endocytic mechanism, which might be a form of CIE. In good agreement, to date no clathrin-coated vesicles have been identified at the parasite surface and a previous study did not implicate CHC in endocytosis in *T. gondii* (21).

Next, we determined the average time required for capping under the same conditions as above (Fig. 1G). In control parasites, capping occurs within 26.4 ± 7.3 seconds. Interference of F-actin using CD did not have a significant effect on capping time (28.9 ± 22.4s). In contrast, interfering with secretion of micronemes by expression of DrpB-DN resulted in a significantly longer capping time of 62.5 ± 36.0 seconds (in the few instances when capping could be observed). Incubation of parasites with endocytosis inhibitors (Phenylarsine or Trifluorperasine) resulted in a complete block of capping. No difference was observed upon expression of CHC-Hub (-Shield: 30.8 ± 17.8 s vs +Shield: 35.1 ± 14.2 s), again suggesting CIE.

When motility was analysed using the same conditions, we found a clear correlation between capping and motility. We confirmed previous findings, demonstrating that interference with the acto-myosin system of the parasite results in significantly reduced overall gliding motility (Fig.1H, I). Importantly, parasites still capable of gliding did so at similar speeds as control parasites (Fig. 1J), as described previously (7). In contrast, conditions that resulted in less and/or slower capping resulted in both fewer parasites capable of initiating gliding motility (Fig.1H, I), and parasites moving significantly slower (Fig.1J), when compared to controls.

Together these data suggest a link between parasite motility and capping, and that the parasite is capable of ensuring membrane balance by a secretory-endocytic cycle, as proposed by the fountain-flow model (Fig.1A).

### Extracellular *T. gondii* tachyzoites are capable of endocytosis

While it has been recently demonstrated that intracellular parasites are capable of taking up host proteins by an endocytosis-like mechanism (16), to date the presence of endocytic activity of extracellular parasites is still under debate. In order to visualise membrane turnover, we assessed the capacity of *T. gondii* to take up fluorescent lipids, such as Lysophosphatidyl choline (LPC), Lysophosphatidic acid (LPA), or BODIPY (BP) (Figure 2A). In all cases, we observed efficient uptake of the lipids, characterised by the occurrence of discernible green fluorescent vesicles inside the parasite, that might be associated with the parasite’s secretory system (Figure 2A, B). The lipids were taken up at comparable rates (BP:82.9±10%, LPC:66.1±2%, LPA: 67.9±2.3%) and uptake only occurred when parasites were incubated at 37°C, while no uptake was obvious at 4°C or when dead parasites were incubated with these lipids (Fig. 2A, B, S1). Together these data demonstrate an active uptake of phospholipids in the majority of extracellular parasites.

**Figure 2:**
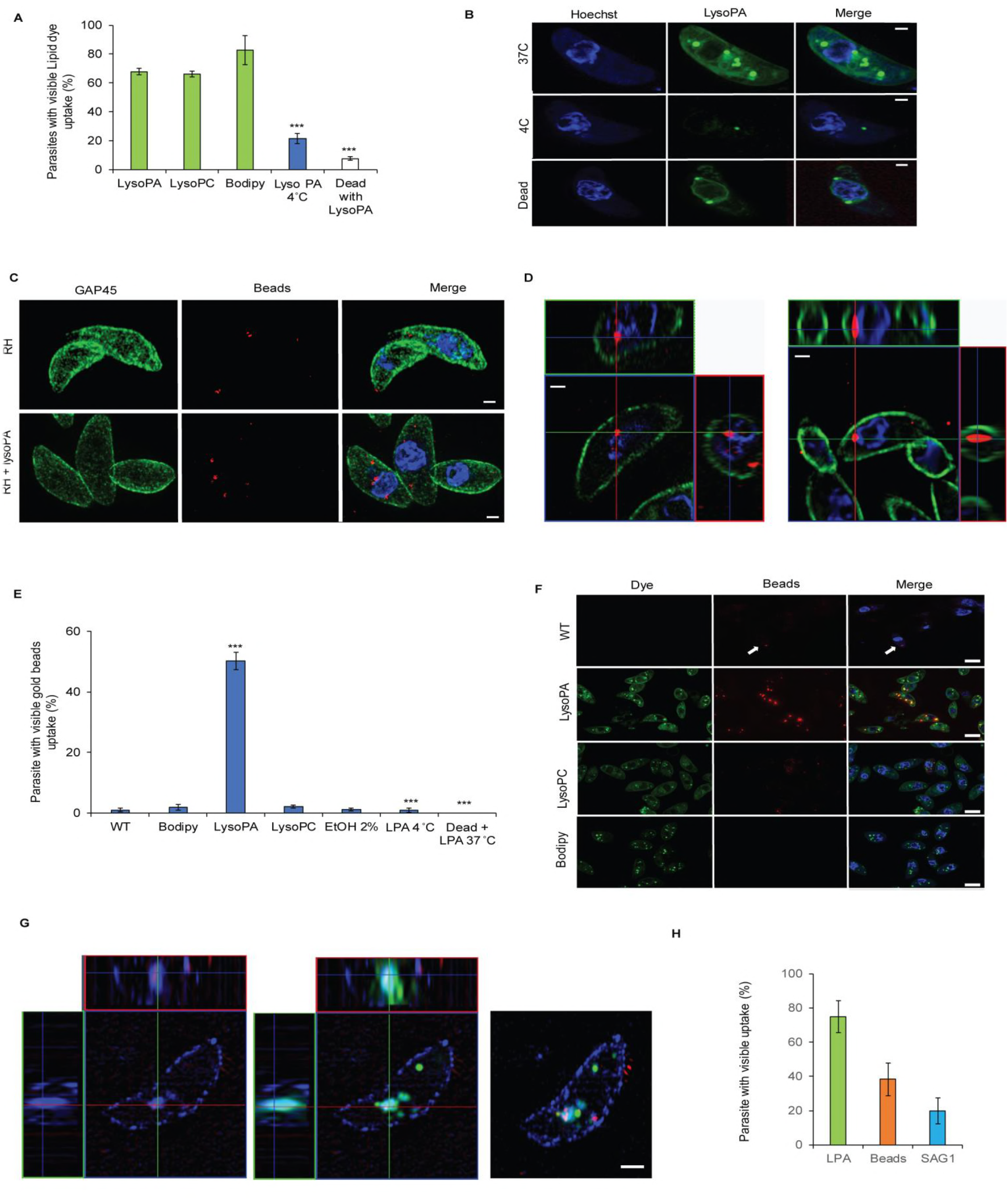
Phospholipids and nanogold particles are incorporated by extracellular tachyzoites. (A-B): Uptake of phospholipids: lysophosphatidic acid (LysoPA), lysophosphatidic choline (LysoPC) and Bodipy were analysed at 37°C and 4°C. Pre-fixed parasites with PFA were also incubated with LPA at 37°C. Incubation at 37°C demonstrates the uptake of all tested molecules. Uptake is inhibited at 4°C or when the parasites were fixed prior to incubation. (A) Mean values of three independent assays are shown ± SEM. *** *p*-value <0.001 in a two-tailed Student’s t-test. (B) Examples of images obtained for the quantification as shown in (A). Scale bar 1µm. (C-D): Uptake of nano gold particles (NGP) was tested with 10 µm Cy5-gold beads. Parasites were imaged using 3D-SIM microscopy. NGP were found to accumulate below GAP45 as illustrated by either maximum intensity projection (C) or ortho-view (D). Scale bar 1µm. (E-F): NGP uptake is significantly stimulated when parasites were co-incubated with LPA. Quantification (E) of the NGP uptake under different conditions: control (WT), or in presence of Bodidpy, LPA, LPC, or EtOH revealed that LPA is a strong stimulator of uptake, as illustrated in an IFA-overview in (F). Scale bar 5µm. Mean values of three independent assays are shown ± SEM. *** *p*-value <0.001 in a two-tailed Student’s t-test. (G-H): Uptake of membrane material was determined by co-incubating RH parasites with an antibody against sag-1 (*blue*), LPA (*green*), and NGP (*red*). Parasites were imaged by 3D-SIM microscopy ((G), Scale bar 1µm), and the uptake quantified (H). Mean values of three independent assays are shown ± SEM.

Next, we assessed if the observed uptake is due to an endocytosis-like mechanism. In the absence of well-established protein markers for endocytosis, we decided to analyse the uptake of 10 nm nanogold particles (NGP) that are regularly used to analyse endocytosis in other eukaryotes (32). When wildtype parasites were incubated with NGP it was possible to detect NGP in vesicular structures inside the parasite (Fig.2C, D, S1, S2, Video S4). Interestingly, we found that NGP uptake is significantly stimulated when parasites were incubated with LPA, but not LPC or BODIPY, with up 50% of the parasite population demonstrating NGP uptake in presence of LPA, compared to less than 5% in its absence (Fig.2E, F, S2). NGP uptake was inhibited by incubation at 4°C or when dead parasites were analysed (Fig.2A), confirming that this represents active uptake rather than passive diffusion of NGP. LPA is a phospholipid that is naturally present in serum (33) and has been reported to be an endocytosis stimulator in mammalian cells (34, 35). No toxic effect or alteration in invasion, replication, or parasite morphology was observed in parasites incubated with LPA (Figure S2B, C, D) or LPC (data not shown). Triggering microneme secretion by addition of 2% ethanol did not lead to any significant increase in NGP uptake (1.1±0.4%, Figure 2E).

We hypothesised that NGP uptake is an endocytic process that can be triggered by LPA. Consequently, we speculated that parasite surface proteins, such as the GPI-anchored surface antigen 1 (SAG1), are likely to be recycled. To test this hypothesis, we incubated parasites in the presence of LPA and α-SAG1. After 30 minutes, parasites were fixed, stripped of unbound α-SAG1, and analysed for uptake of the antibody. We found that treatment of parasites with LPA leads to internalisation of α-SAG1 in ∼20% of all parasites (Fig.2G, H), demonstrating the existence of an endocytic recycling pathway. Importantly, internalised α-SAG1 co-localises with LPA and internalised NGP, demonstrating that α-SAG1, LPA, and NGP utilise the same pathway. However, we can only speculate as to why the observed percentage of parasites recycling α-SAG1 is significantly lower compared to NGP uptake. It is possible that SAG1 is rapidly recycled or processed in the VAC (16).

In contrast to proteinaceous markers such as α-SAG1, the use of a non-reactive particle like NGP therefore allowed us to detect endocytosis more efficiently, and so we chose to continue this study using LPA and NGP to further characterise uptake dynamics and the uptake pathway.

### Dynamics of the endocytic cycle

Next, we analysed uptake of LPA and NGP over time (Fig. 3A, B, S3). Interestingly, intracellular LPA can be detected rapidly (from 17.0±1.6% 1 min after to 67.9±2.3% 30min after addition), while NGP detection lagged behind (0.3±0.3% 1min after to 50.3±2.8 30min after addition, Fig.3A). Though this difference could be explained by a weak signal to noise ratio of single gold beads, requiring accumulation of NGP in endocytic compartments before they can be detected, it is also possible that this lag phase is due to the requirement of LPA to stimulate endocytosis of more bulky material, as described in other eukaryotes (34, 35). Furthermore, the average number of intracellular vesicles increases over time, from ∼2 at 5 minutes to ∼5 at 30 minutes after addition of LPA (Fig. 3B, S3).

**Figure 3:**
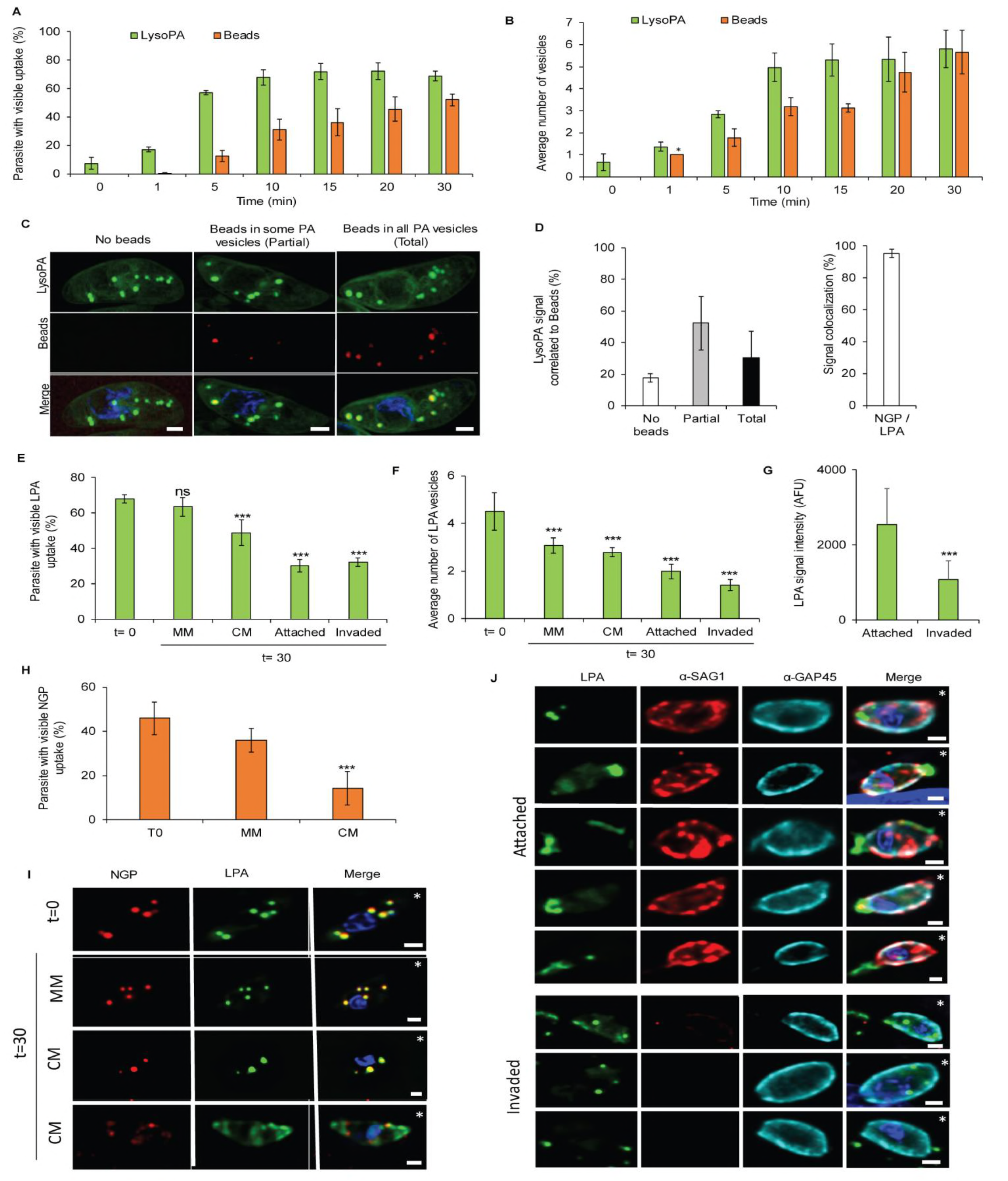
LPA and NGP uptake is part of an endocytic-exocytic cycle. (A-B): Dynamics of uptake were analysed by time point assay from 0 to 30 minutes after addition of LPA and NGP. Average number of parasites that show overall uptake (A) and average number of vesicles per positive parasite (B) was determined. Mean values of three independent assays are shown ± SEM. (C-D): An)alysis of LPA and NGP uptake at 30min. Three different observations were made (C): Parasite with LPA vesicles without NGP (No beads), NGP in some LPA vesicles (Partial) and NGP in all LPA vesicles (Total), Scale bar: 1 µm: Each phenotype was quantified and the colocalization ratio between NGP and LPA calculated (D). Mean values of three independent assays are shown ± SEM. (E-J): Exocytosis of LPA and NGP was evaluated 30 minutes after placing parasites under three conditions: Minimal media (MM), complete media (CM) and complete media with host cell. In the latter case, invaded and attached parasites were analysed separately. A clear diminishing of the signal is observed over time in stimulating conditions, demonstrating secretion of previously endocytosed material. This was seen for both NGP and LPA. Percentage of LPA positive parasites (E) and average number of vesicles (F) was calculated. Mean values of three independent assays are shown ± SEM. *** *p*-value <0.001 in a two-tailed Student’s t-test. (G): Average vesicle signal intensity was calculated between attached and invaded parasites, n=25. (H): Comparison of the percentage of NGP positive parasites transferred to minimal (MM) and complete media (CM). Mean values of three independent assays are shown ± SEM. *** *p*-value <0.001 in a two-tailed Student’s t-test. (I): Representative images of parasites transferred to minimal or complete media. Scale bar 1µm. * Indicate the apical pole of the parasite (J) Representative images of parasites transferred onto host cells. Sag-1 staining (prior to permeabilisation) was used to differentiate intra-from extra-cellular parasites. Scale bar 1µm. * Indicate the apical pole of the parasite

Next we analysed the location of LPA-and NGP-positive vesicular structures inside the parasite (Fig.3C, D). We observed three different distributions (Fig. 3C, D): 1) parasites with LPA-positive vesicles without NGP (“No beads”, 17.5±2.6%); 2) parasites with LPA-positive vesicles where some of the vesicles also contain NGP (“Partial”, 52.1±16.9%); and 3) parasites where all LPA-positive vesicles contain NGP (“Total”, 30.2±16.9%). These data correlate well with the dynamic of uptake described above, since LPA uptake itself appears to occur earlier than NGP and therefore more LPA-positive structures are expected. Importantly, in most cases (95.2±2.5%, Fig. 3D) we observed that NGP-positive vesicles are also LPA-positive, demonstrating that they converge in the same trafficking pathway.

If a secretory-endocytic cycle operates within the parasite, it is possible that material that entered via the endocytic route could be recycled and secreted. To test this hypothesis, parasites were pre-treated with LPA and NGP for 30 min, before excess material was washed away, and parasites transferred to new dishes containing minimal media, complete media, or host cells with complete media for 30 min before fixation. We found that NGP and LPA signal changed when parasites were placed into complete media or dishes containing host cells (Fig.3E, F, G). In minimal media, the percentage of positive parasites was similar prior to washing, for both LPA (68.0±3.7% versus 58.9±3.4%) and NGP (42±2.0% versus 33.6±4.3%). In contrast, when the parasites were placed in complete media or in the presence of host cells, a drastic reduction in signal was detected for both LPA (67.0±2.3% vs 32.0±2.4%, Fig. 3E) and NGP (42.9±2.0% vs 8,9±2.8%, Fig 3H). Importantly under these conditions a different distribution of vesicular structures became evident (Figure 3I, J), where LPA was observed at the apical and basal ends of the parasite, as well as at the parasite surface. LPA was also observed in trails when parasites were incubated with host cells, suggesting secretion of the LPA/NGP-positive vesicles. This is supported by a decreased intensity of LPA signal between attached and invaded parasites (Fig. 3G). Taken together, these data indicate that a complete endocytic-exocytic cycle is occurring under these conditions.

### LPA and NGP co-localise inside vesicles along the secretory pathway of the parasite

We were interested in defining the pathway followed by endocytosed material in detail. To this end, we performed co-localisation assays of LPA-positive vesicular structures with previously described markers of the parasite secretory pathway (Table 1, Fig. 4A, B). The highest accumulation of LPA could be observed in the VAC (30.7±8.6%), a plant-like vacuole in the parasite (26, 27). The second highest colocalization was observed with RAB18 (26.6±3.72) a marker of the ER (22). Less accumulation could be seen with other organelles, such as endosomes (VPS53: 12.1±5.1%), Golgi (Rab4: 8.7±3.4% and CHC: 9.3±2.9%), rhoptries (ROP1: 9.8±3.2%), and endosome-like compartment (ELC) (pro-M2AP 3.3±0.8%). No colocalization was observed with RAB2 (ER), VPS35 (ELC), MIC2 (micronemes), and GRA1 (dense granules; Fig. S4). Together, these data demonstrate that exogenous material is taken up via a conserved endocytic pathway, following previously identified secretory and endocytic organelles in the parasite.

**Figure 4:**
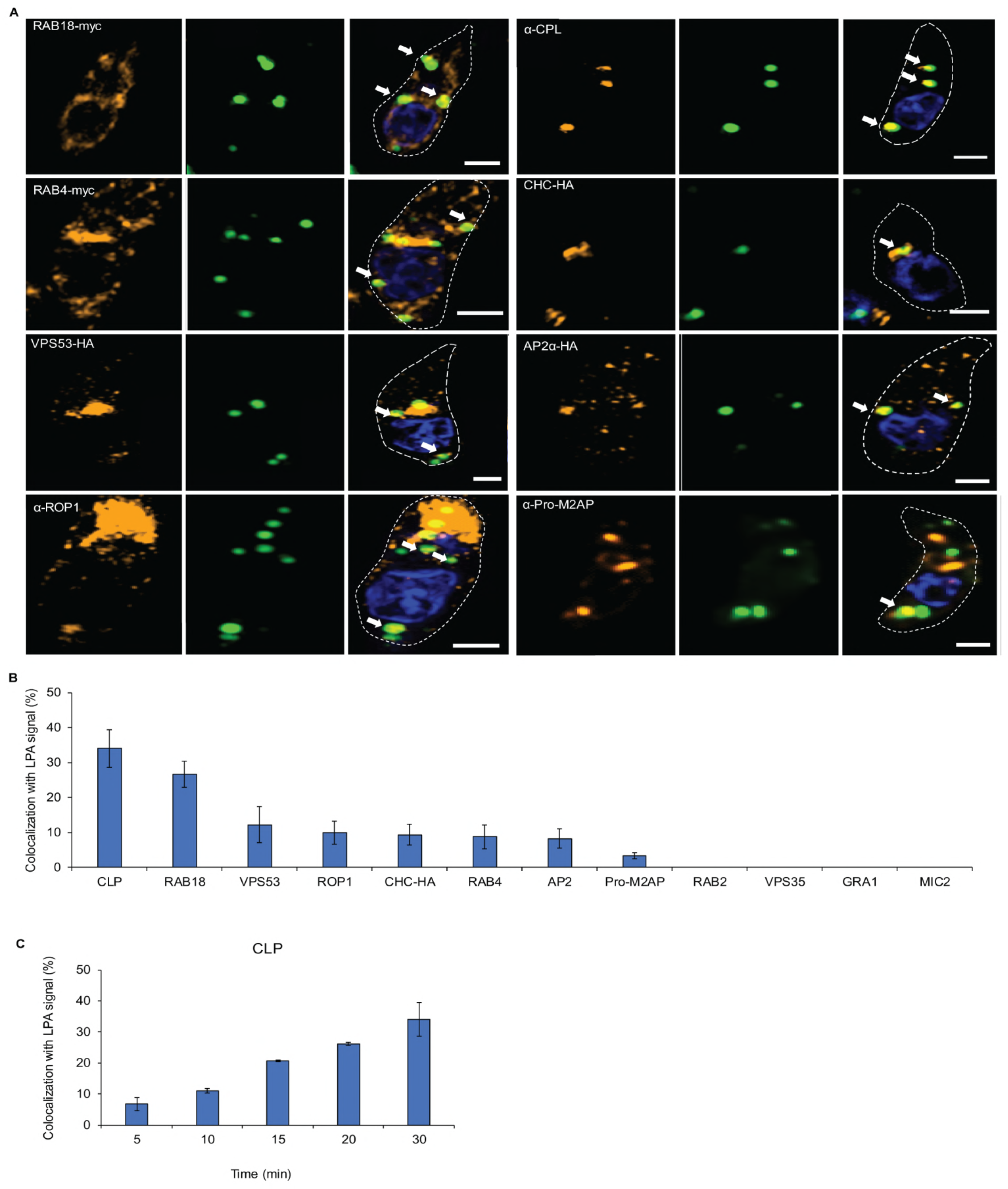
Endocytosed material traverse the secretory system. (A-B): LPA vesicle localisation was determined using indicated parasite strains expressing markers for the parasites trafficking system: dd-RAB18-myc: ER, dd-RAB4-myc: Golgi, VPS53-HA: TGN, VPS35-HA: Retromer/ELC, CHC-HA: Golgi, TGN, AP2α-HA parasites (21, 22) (63, 67). Antibodies against the VAC (α-CPL, (27)), rhoptries (α-ROP1), micronemes (α-MIC2), and dense granules (α-GRA1) were used after fixation on RH parasites. (A): Representative pictures of colocalizations are shown (see also FigureS4), Scale bar 1µm. (B): Quantification of colocalizations. Mean values of three independent assays are shown ± SEM. (C): LPA accumulates in the VAC over time. Percentage of colocalization of LPA and CPL, was determined from 0 to 30min after LPA addition. Mean values of three independent assays are shown ± SEM.

To investigate the nature of the identified compartments and the origin of uptake in more detail, we performed correlative light and electron microscopy (CLEM). Parasites were incubated with LPA and NGP for 30 minutes before fixation (Fig. 5A). At least two types of vesicles could be identified: vesicles with a similar density as the cytoplasm (Fig. 5A 1 and 2) and other more translucent vesicles (Fig. 5A 3). Next, classical Transmission Electronic Microscopy (TEM) was used (Fig. 5B, C, D, E, G). As observed by IFA and CLEM, NGP were found in different locations within extracellular tachyzoites. NGP accumulated with different densities inside vesicles (Fig.5B, C) and were found in at least 3 types of vesicles: large translucent vesicles (300nm - 500nm, Fig. 5C panel 1), medium size dense vesicles (215nm-375nm, Fig.5C2) and small vesicles (80nm-200nm, Fig. 5C panel 3 and 3’). In good agreement with the co-localisation experiments, NGP were frequently observed inside the VAC (Fig. 5D). In some instances, NGP were also detected inside the rhoptry bulb both in TEM and Tomography, confirming the co-localisation with ROP1 observed by IFA, and suggesting that some of the vesicles could fuse or exchange material with the rhoptries (Fig. 5E, F). In contrast, NGP were never seen in dense granules or micronemes. We also identified invaginations at the surface of the parasite that contain NGP, probably representing the point of uptake. These structures are delineated by the plasma membrane, demonstrating that NGP are actively taken up in an endocytic-like process. Importantly, these invaginations are not electron dense and a classical clathrin cage could not be detected, suggesting a clathrin-independent uptake mechanism. (Fig. 5G).

**Figure 5:**
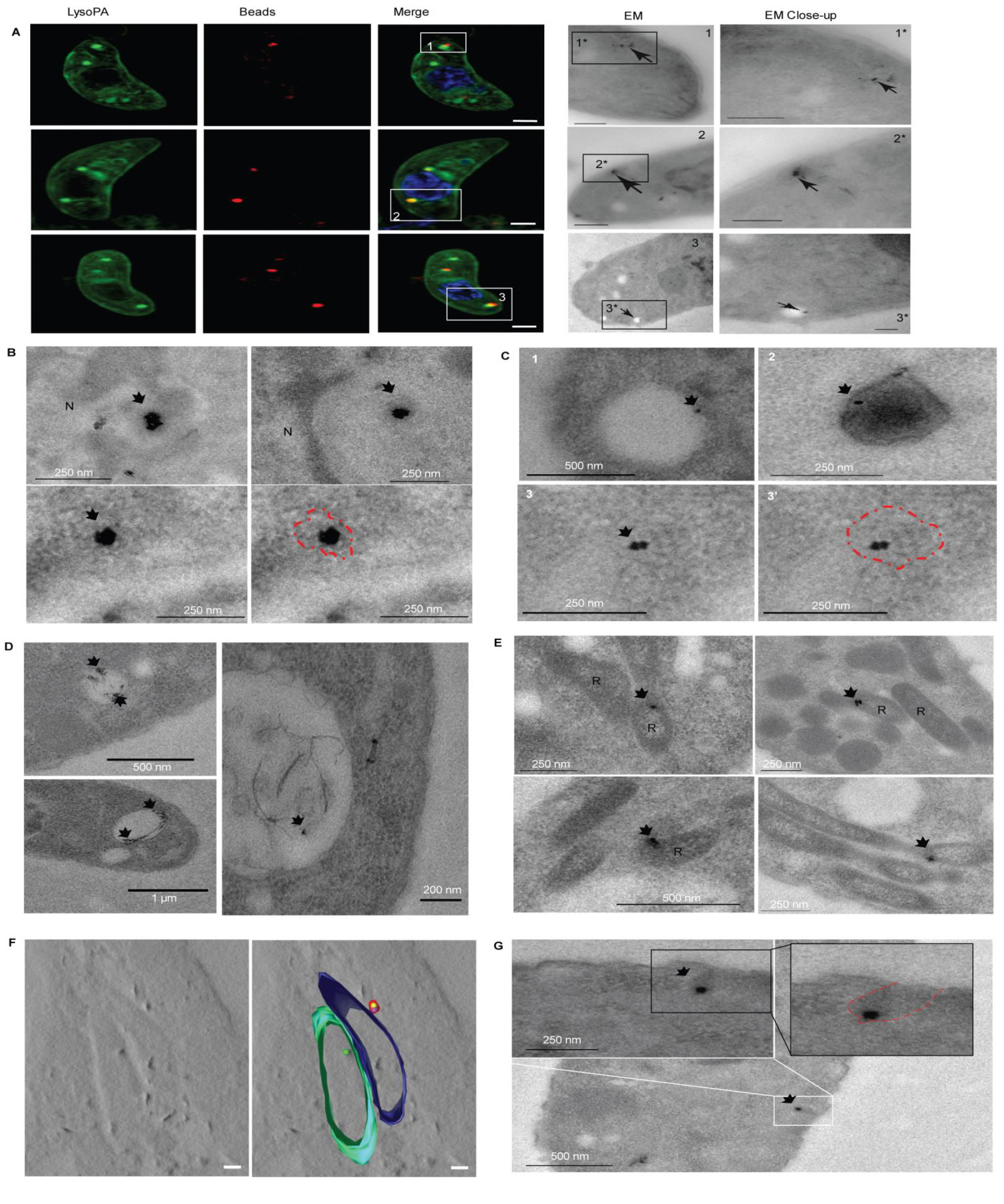
Trafficking of NGP and LPA confirmed by EM. (A): CLEM imaging of LPA and NGP signal. After the uptake experiment parasites were imaged by IFA (Scale bar 1µm) before processing for EM (Scale bar 200nm) to confirm that the signal is inside vesicles of different types and therefore that endocytic uptake has occurred. White square, close up illustrated by the first TEM image; black square, TEM close up. Black arrows indicate the NGP. (B-G): TEM localisation of NGP. Scale bar size is indicated on the images. Black arrows indicate the NGP. For the less dense vesicles and membrane invagination a picture where the delimitation is highlighted in red is present (B): Bead aggregation observed inside vesicles. Parasite nucleus is labelled with N. (C): Representative images of the different types of vesicles observed: 1) large translucent vesicles (300nm - 500nm), 2) medium size dense vesicles (215nm-375nm) and 3,3’) small vesicles (80nm-200nm) (D): Localisation of the NGP in the VAC, (E-F): Localisation of the NGP in rhoptry bulbs (Labelled with R) by both TEM (E) and tomography (F). (G): Image of a potential entry point as an invagination of the plasma membrane containing NGP.

### Endocytosis is linked to motility of *T. gondii* tachyzoites

As described above (Fig. 1), we were able to observe a link between capping of beads and motility. To determine if a similar correlation between retrograde membrane flow and endocytosis exists, we co-incubated parasites with 40nm beads (used in the capping assay) and LPA (Fig. 6A, B, C). The addition of lipids did not significantly increase overall capping activity of *T. gondii* (Capped-LPA: 30.6±5.8, Capped +LPA 33.0±6.1). Interestingly, we observed that almost all capped parasites (85.6 ± 9.4%) also presented LPA positive vesicles, as illustrated in Figure 6B, while parasites that only bound latex beads to their surface did not show a high percentage of LPA uptake (11,7 ± 8.4%, Fig. 6A, B). This correlation between capping and LPA uptake was also observed using live microscopy (Fig. 6C, Video S5).

**Figure 6:**
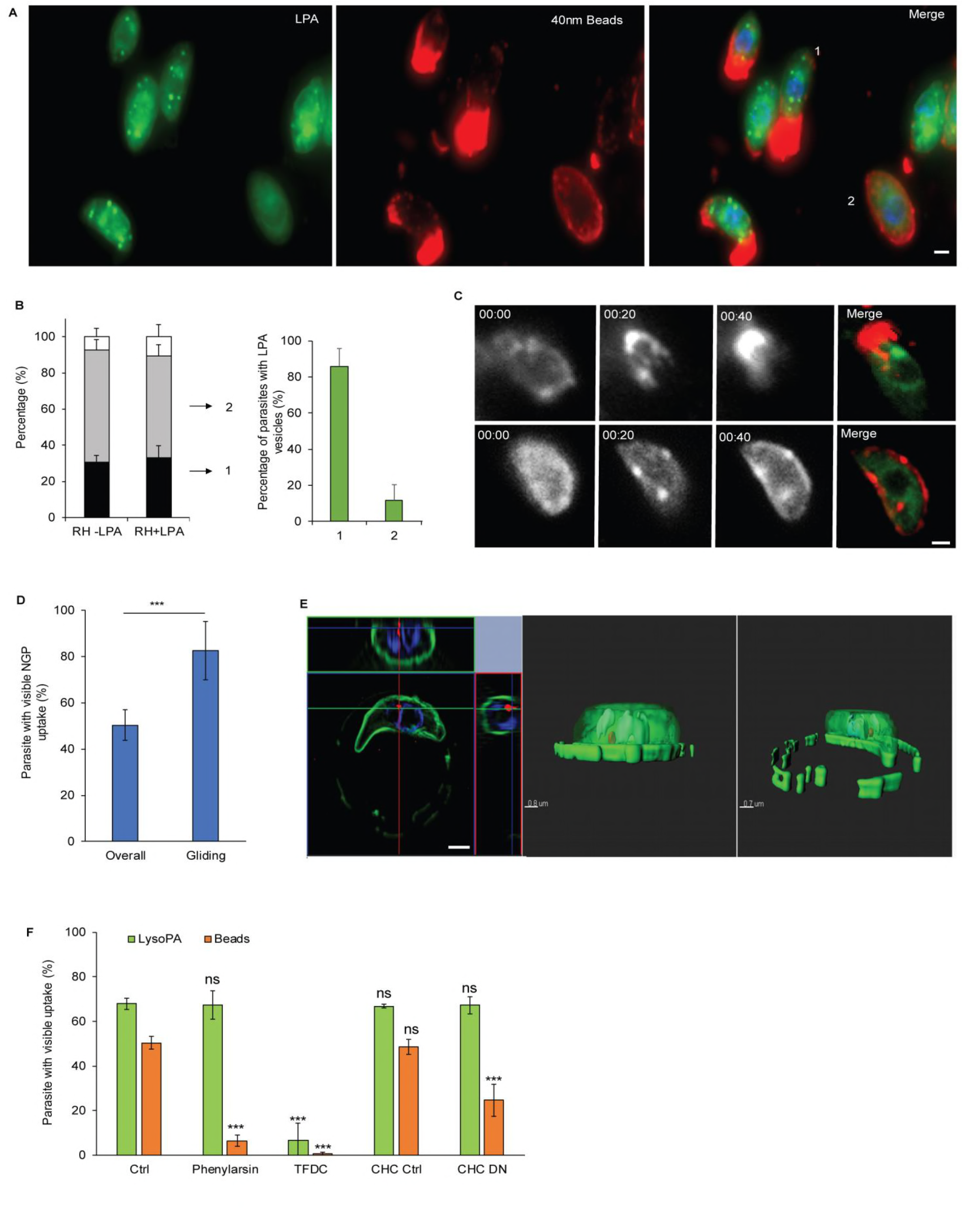
Correlation between endocytosis, retrograde flow, and motility. (A-C): Association of capping and endocytosis was analysing by adding LPA during the bead translocation experiment. (A): Representative picture of capping and LPA uptake. An example of each phenotype can be observed: capped (1), bound (2). Scale bar 1µm. (B): Quantification of the percentage of capped (1), bound (2) and unbound parasites that also took up LPA. Note that “unbound” represents parasites, where beads did not bind and so no statement regarding occurrence of retrograde membrane flow is possible. Importantly, most parasites which translocated beads also endocytosed LPA. Mean values of three independent assays are shown ± SEM. (C): Time lapse analysis of bead translocation in the presence of LPA. An example of a parasite, where translocation occurred (top) and did not occur (bottom) is shown. Scale bar 1µm. (D-E): Correlation between gliding and endocytosis was analysed by adding LPA and NGP during the motility assay. (D): Quantification of NGP uptake between the overall population of parasites and gliding parasites. Mean values of three independent assays are shown ± SEM. *** *p*-value <0.001 in a two-tailed Student’s t-test. (E): 3D-SIM microscopy ortho-view of a representative parasite (Scale bar 1µm) and 3D reconstructions images. (F): Effect of inhibitors and clathrin DN on the LPA and NGP uptake; both endocytosis inhibitors, which efficiently inhibit retrograde flow and gliding motility, inhibit the uptake. Expression of dominant negative CHC, which did not significantly impact motility, appears to have a mild effect on NGP (but not LPA) uptake.

To address the importance of endocytosis during gliding motility, parasites were incubated with LPA and NGP and put on coverslips coated with FBS to perform gliding assays. After fixation, parasites were incubated with α-SAG1 antibody to observe trail depositions (Fig. 6D, E, Video S6). We obtained a similar percentage of NGP uptake as in our previous experiments (50.3±6.7%). Strikingly, considering only gliding parasites, i.e. those that deposited trails, NGP uptake markedly increased (82.5±12.5%).

Finally, we used different mutant strains and drugs as above (Fig.1, 6F). Endocytosis inhibitors that impacted both gliding and capping strongly diminished NGP uptake. In addition, phenylarsine did so without impacting LPA uptake (LPA: 67.3±6.2%, NGP: 6.5±2.4%), whereas TFDC inhibited both LPA and NGP uptake (LPA: 6.8±7.4%, NGP: 0.6±0.6%). We also tested the DN CHC strain, which does not significantly affect either gliding or capping. We observed a slightly reduced uptake of NGP (Ctrl: 48.5±7.1% vs DN:24.6±1.7), whereas no reduction in LPA uptake was observed. This suggests that CHC is not critical for endocytosis and that the observed reduction of NGP uptake might be due to downstream effects caused by blocking post-Golgi transport through CHC-DN expression (21).

Taken together, these results demonstrate a clear link between retrograde membrane flow, endocytosis, and parasite gliding motility, and therefore fully support the fountain flow model.

## Discussion

### A link between gliding motility, retrograde membrane flow, secretion, and endocytosis

Apicomplexan gliding motility has been considered a “unique” form of cell motility that depends on special secretory organelles found in apicomplexans, which secrete transmembrane proteins to link the acto-myosin system anchored in the IMC of the parasite to the substrate during motility. The unusual features of parasite actin, combined with the absence of direct homologues of many classical actin binding partners, led to the hypothesis that apicomplexan motility is distinct from other classical eukaryotic mechanisms. Here we argue that this form of motility is similar to amoeboid motility and that many of the identified crucial elements for gliding motility play analogous roles compared to factors found in other eukaryotes. A good example are micronemal proteins, which act similar to integrins and often contain integrin-like extracellular domains, such as MIC2 (36). Micronemal proteins are required for attachment to the surface and have been recently shown to interact with F-actin via the glideosome-associated connector (GAC) (37), which appears to fulfil an analogous role to talin, which connects integrins to F-actin in motile eukaryotic cells (38). Interestingly, and very similar to findings in apicomplexan parasites, recent results in motile eukaryotic cells have led to a reassessment of the traditional motility models. Of note, analysis of cell migration in a 3D-environment, which is arguably more physiological than a 2D-environment, has led to a rich body of literature describing new motility mechanisms such as blebbing, osmotic engines, fountain-flow, membrane stretch, etc. (10, 39-43). Although lengthy description of these various models is outside the scope of this discussion, it is important to note that motile cells can move by employing more than one mechanism depending on the environmental conditions encountered.

In the case of apicomplexan parasites, the current model describes a simple linear motor that is required for all motility processes of the parasite, including host cell invasion (29). However, the parasite presumably encounters 2D, 3D, constricted, adhesive, and non-adhesive environments during its life cycle, requiring adaptations to the respective surrounding, just as other eukaryotic cells, where different forms of motility have been described. Importantly, recent studies demonstrate that the linear motor model cannot explain all forms of motility. This includes 1) Biophysical studies on malaria sporozoites demonstrating the discrete, localised turnover of attachment sites that are not evenly translocated along the surface of the parasite (6). 2) Different types of motility observed in *Eimeria* sporozoites (bending and pivoting). While pivoting appeared substrate and actin dependent, bending appeared resistant to the actin disrupting drug Cytochalasin B (44). 3) Reverse genetic approaches demonstrated that parasites remain motile in the absence of key molecules of the linear motor machinery, including parasite actin (8, 45-48). 4) Parasite F-actin dynamics are complex, and actin is not only localised (as predicted by the linear motor model) between the IMC and the plasma membrane of the parasite, but is found in other areas of the parasite as well (49). 5) Retrograde membrane flow of malaria sporozoites occurs even at relatively high concentrations of the actin-disrupting drug CD (11) and in the case of *T. gondii*, occurs upon disruption of crucial elements of the parasite acto-myosin system (7).

Importantly, retrograde membrane flow is implicated in many different motility modes of eukaryotic cells and recent evidence demonstrates important roles for membrane trafficking in the regulation of cell migration in a variety of contexts. For example, one critical role of an endocytic-secretory cycle is the internalisation and recycling of adhesion receptors, such as integrins or syndecans (50). Another critical role is the maintenance of a constant cell surface (51, 52). Indeed, recent studies suggested the operation of a so-called fountain-flow model that predicts that surface area regulation and retrograde membrane flow is caused by a secretory-endocytic cycle (10). Here we demonstrated that the situation in the apicomplexan parasite is very similar and fully supports the fountain-flow model proposed previously for other eukaryotic cells (53). Previous studies implicated the generation of a retrograde membrane flow in parasite motility (7, 11, 54). Interestingly, membrane flow itself is relatively resistant to actin-modulating drugs, whereas force generation depends on a functional acto-myosin system. Here, we extended our analysis of retrograde membrane flow using a bead translocation assay (7) and demonstrate a strong link between the occurrence of bead translocation, parasite motility, and the operation of an endocytic-secretory cycle. We verified that interference with the acto-myosin system of the parasite has little effect on bead translocation, while blocking apical microneme secretion or endocytosis abrogates bead translocation and parasite motility (Fig. 1F, G, H, I, J)

### Extracellular parasites can take up exogenous material and recycle surface proteins

Uptake of host cell material from intracellular parasites has recently been demonstrated and it appears that this material is taken up by an endocytic route that merges with the secretory pathway of the parasite (15). This endocytic process appears to occur rapidly, with endocytosed proteins eventually reaching the vacuolar-like compartment (26, 27) of the parasite, in which they are digested (16). However, it was previously unclear whether extracellular parasites are capable of similar uptake, especially since a number of trafficking factors normally involved in endocytosis have been shown to be required for the transport of material to unique secretory organelles, leading to the hypothesis that endocytic factors have been repurposed during apicomplexan evolution (55). Here we demonstrate that extracellular parasites, like intracellular parasites, are capable of endocytosis, and that the parasite appears to recycle surface proteins, such as SAG1. Interestingly, we observe that lipid dyes are efficiently taken up, but only LPA appears to stimulate endocytosis of larger material, such as 10nm gold beads. It is thus likely that LPA triggers endocytosis via an as yet unknown signalling cascade. LPA has been demonstrated in numerous systems to activate endocytic uptake (34, 35), and can be converted to phosphatidic acid (PA) (56), which can act as a second messenger. For example it can promote both CME and CIE (57, 58), and can also lead to the recruitment and activation of proteins directly involved in the trafficking machinery (59). Interestingly, phosphatidic acid (PA) has been previously implicated in regulating microneme secretion and invasion in apicomplexan parasites (60), making it attractive to speculate that PA is a central mediator for the secretory-endocytic cycle. Stimulation of parasites with LPA leads to efficient uptake of NGP, which co-localise with LPA-positive vesicular structures. Importantly, we found that these structures co-localise with markers of the endomembrane system of the parasite, such as the ER, Golgi, VAC, and rhoptries, with a clear accumulation in the VAC, suggesting that the same system is employed as for the uptake of host cell material during intracellular stages (16). Ultrastructural analysis confirms these findings and suggests that the material is taken up at surface invaginations that appear to be distinct from the micropore of the parasite (61). To date, our attempts to obtain mechanistic insight regarding the endocytic process have been unsuccessful. While established endocytosis inhibitors show the expected effects, i.e. marked decrease in uptake of material, analysis of dominant negative mutants for clathrin or dynamins had no significant effect on endocytosis. Furthermore, no clathrin-coated pits, a hallmark of CME, could be detected in our ultrastructural analysis. While this might suggest that the analysed factors are not involved in endocytosis, it is also possible that the conditional mutants used do not have the right kinetics for downregulation of the respective genes, since this might already cause parasite death within the host cell, before extracellular endocytosis can be analysed. Therefore, faster conditional regulation systems should be used in future studies to re-analyse these factors, such as the auxin-inducible degron system (62).

We demonstrate here that *T. gondii* is capable of performing endocytosis in the extracellular stage by presenting a method to visualise internalization and recycling of both surface proteins and exogenous material (SAG1 and LPA/NGP, respectively). It is interesting that NGP were found in the rhoptries, but not micronemes, since components of both organelles are delivered by a common secretory system (63). One possibility is that NGP accumulate in the rhoptries, since these are not secreted in extracellular parasites, while micronemes can be secreted (for example during motility). Indeed, we demonstrate here that the NGP are secreted upon incubation of parasites in complete media or on host cells, providing evidence for the existence of an endocytic-recycling pathway that operates in stimulated parasites.

### The Fountain-Flow-Model for apicomplexan parasites

Based on these data, and previous data that suggested an important role of the acto-myosin-system as a clutch for the transmission of force (7), we suggest that the generation of retrograde membrane flow is critical for gliding motility. This flow can be generated by the acto-myosin-system of the parasite as well as regulated secretion and endocytosis, as suggested for other eukaryotes (53). According to this model, apical microneme secretion needs to be balanced by endocytic recycling, resulting in a lipid drive or fountain flow (10). In good agreement with this model (Figure 7), we demonstrate that gliding parasites generate retrograde membrane flow while simultaneously internalizing NGP; furthermore, inhibitors of uptake also block retrograde membrane flow and gliding. In contrast, interference with the acto-myosin system does not cause a block in retrograde membrane flow or uptake, resulting in parasites that glide less, which can be attributed to less surface attachment, as previously shown (7, 8)

**Figure 7:**
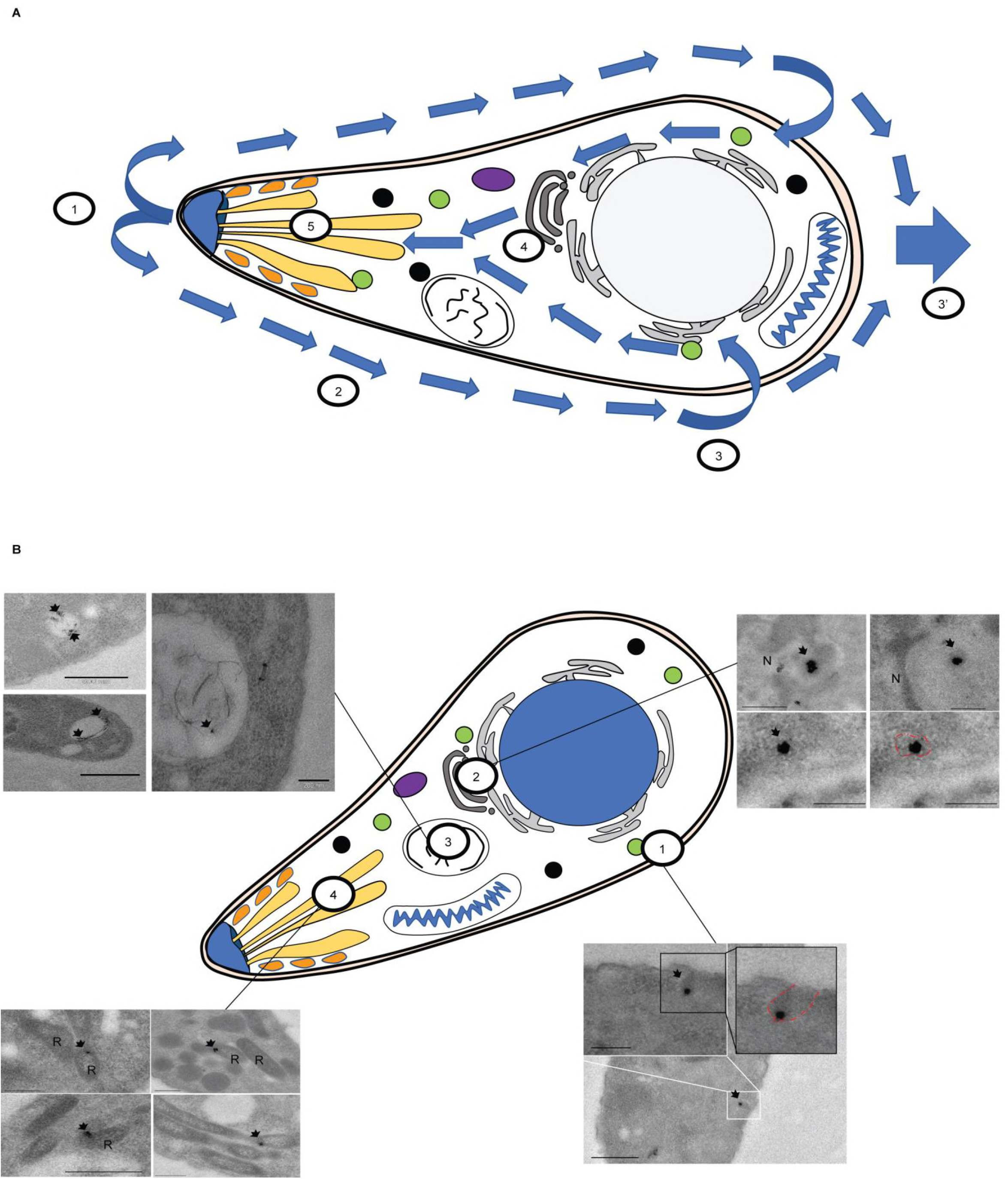
Fountain flow model applied to *T. gondii*. (A): Model of the fountain flow applied to *T. gondii*. 1: Secretion from the secretory organelles at the apical end. 2: Retrograde flow allows bead translocation along the plasma membrane. 3 and 3’: After translocation, membrane material can be either recycled (3) or left in a trail (3’). 4: Endocytosed material is trafficked inside the parasite along the secretory system and accumulates in the VAC. 5: Incorporated material can be secreted in a complete endocytic-secretory cycle. (B): Summary of the different locations observed in TEM. 1: Entry from the plasma membrane via an invagination. 2: Perinuclear localisation, likely to be ER or Golgi. 3: Trafficking to the VAC. 4: Trafficking to the secretory organelles (Rhoptry).

### Outlook and Summary

While this study reconciles many previously conflicting data, it does not directly address the mechanisms involved in endocytic uptake, as discussed above. It will now be important to identify the crucial trafficking factors directly involved in endocytosis. It is likely that the parasite evolved specific structures for endocytic uptake, as suggested by the presence of the invagination seen in TEM analysis. A recent genome wide screen (64) demonstrated the essentiality of hundreds of hypothetical genes. Many of them might well be involved in an essential uptake pathway, required for gliding motility and host cell invasion. Therefore, with the establishment of a reliable uptake assay, it is now possible to phenotypically screen for essential genes involved in this process, which will not only result in a fundamental understanding of this process, but also in the identification of novel intervention strategies.

### Materials and Methods

#### Cloning DNA constructs

All primers used in this study are listed in Table 2 and were synthesised from Eurofins (UK).

#### Culturing of parasites and host cells

Human foreskin fibroblasts (HFFs) were grown on TC-treated plastics and maintained in Dulbecco’s modified Eagle’s medium (DMEM) supplemented with 10 % Foetal Bovine serum, 2 mM L-glutamine and 25 mg/ml gentamycin. Parasites were cultured on HFFs and maintained at 37 °C and 5 % CO_2_.

#### *T. gondii* transfection and selection

To generate stable parasite lines, 1×10^7^ freshly lysed RH *Δhxgprt* or RH-DiCre *Δku80* parasites were transfected with 20 µg of DNA by AMAXA electroporation. Drug selection was carried out with either mycophenolic acid and xanthine as described in (65), or with bleomycin.

### Generation of parasite lines

The following strains have been previously produced and published: CHC-HA, dd-DrpB_DN_, dd-CHC_DN_, *act1* cKO, *mlc1* cKO, *myoA* KO, dd-Rab18-myc, dd-Rab4-myc, and dd-RAB2-myc (7, 21-23)

#### VPS35-HA, VPS53HA

C-terminal 3xHA epitope endogenous tagging of the *vps35* and *vps53* genes was carried out by the ligation-independent cloning (LIC) strategy as previously described (66). Briefly, 15 μg of each plasmid was linearised by EcoRV (LIC *vps35-HA)* or PstI (LIC *vps53-HA)* within the homologous region for efficient homologous recombination and was transfected into *Δku80* parasites. The resultant transfectants were selected for clonal lines expressing VPS35-HA or VPS53-HA in presence of 25 μg/ml mycophenolic acid and 40 μg/ml xanthine and subsequently cloned by limiting dilution. Specific integration was confirmed by analytical PCR on genomic DNA using primers upstream the homology region inserted in the LIC vector, and a reverse primer binding the LIC HA region (Table 2).

#### Internal tagging of AP2α by transient CRISPR-Cas9 expression

sgRNA plasmids were generated by PCR amplification of the guide RNA into a pU6-DHFR plasmid using Q5 mutagenesis kit and following manufacturer procedures (NEB). RH Δku80Δhxgpt parasites were transiently transfected using an AMAXA 4D Nucleofector (Lonza) with Cas9-YFP, each sgRNA plasmid and reparation template PCR product. Briefly, a total of 10 μg of precipitated plasmid DNA and PCR product and approximately 9 x 10^5^ parasites were resuspended in 20 μl of Buffer P3 (P3 Primary Cell 4D Nucleofector X kit S (32 RCT), Lonza). Parasites were transfected in a multi-well format using a programme FI-158, transferred to an HFF monolayer and fixed at 24 and 48 h post transfection. Parasites were considered transfected if expressing GFP-Cas9 in the nucleus. Checking of the integration was made as described above.

#### Inducing conditional knockdown lines and protein expression

dd-DrpB_DN_, dd-CHC_DN_,, dd-DrpB_DN,,_ dd-Rab18-myc, dd-Rab4-myc and dd-RAB2-myc parasites (Table 1) were grown until the vacuoles were ready to lyse. Shield was added 6 hours prior to the parasites being used for experiments. *act1* cKO, *mlc1* cKO were induced as previously described (7).

### Phenotypic characterisations

#### Trail deposition assay

Gliding assays were performed as described before (7). Briefly, freshly lysed parasites were allowed to glide on FBS-coated glass slides for 30 min before they were fixed with 4 % PFA and stained with α-SAG1 under non-permeabilising conditions. Mean values of three independent experiments +/-SEM were determined. Where drugs were used, parasites were pre-incubated for 10 minutes in the respective concentration before the start of the assay: 0.5 µM Cytochalasin-D (CD) (Sigma), 10 µM phenyl arsine oxide (Sigma) or 50 µM Trifluoperazine dihydrochloride (TFDC) (Sigma). The same concentrations were used in the different assays.

#### 2D motility assay

Time-lapse video microscopy was used to analyse the kinetics over a 2D surface similar as previously described (48). Briefly Ibidi μ-dish^35mm-high^ were coated in 100 % FBS for 2 hours at room temperature. Freshly egressed parasites were added to the dish. Time-lapse videos were taken with a 40X objective at 1 frame per second using a DeltaVision^®^ Core microscope. Analysis was made using ImageJ wrMTrckr tracking plugin. For analysis, 20 parasites were tracked during both helical and circular trails with the corresponding distance travelled, average and maximum speeds determined. Mean values of three independent experiments +/-SD were determined.

#### Invasion assay

For the assay, 5×10^4^ freshly lysed parasites were allowed to invade a confluent layer of HFFs for 1 hour after 30 minutes treatment with or without LPA. Subsequently, five washing steps were performed for removal of extracellular parasites. Cells were then incubated for a further 24 hours before fixation with 4% PFA. Afterwards, parasites were stained with the α-IMC1 antibody (8). The number of vacuoles in 15 fields of view was counted. Mean values of three independent experiments +/-SEM were determined.

#### Capping assays

Capping assays were performed as previously described (7). Briefly, Ibidi live cell dishes (29 mm) were coated with 0.1% poly-L-Lysine for 30 min and washed with MilliQ water. Fluorescent latex beads (FluoSpheres^®^, 0.04 µm, Invitrogen) were diluted at 5 µl in 400 µl of a mixture of Hanks Balanced Salt Solution (HBSS) and HEPES (25 mM; described hereafter as H-H buffer). After a short spin (10 s, 6000 g), the supernatant was recovered and left on ice for 30 min before use. Parasites were harvested, pelleted, and resuspended in cold H-H Buffer to achieve 10^7^ parasites/ml. Parasites were then transferred to poly-L-Lysine coated dishes and left on ice for 10 min. 5 µl of diluted beads were added to 250 µl of H-H buffer and added to the parasites. Immediately, the dish was incubated at 37 °C for 30 min. The experiment was stopped by addition of 2 ml of 4 % PFA and incubated at 4 °C for 10 min. The PFA was washed gently and parasite nuclei stained with Hoechst 0.01 %. For time course analysis, parasites were fixed at different time points after the addition of the beads. For the drug and buffer assays, parasites were incubated for 10 min in the buffer of interest before their incubation on the coated dish. In these cases, all other experimental components were also diluted using the same buffer. For each experiment (n), an average of 1000 parasites were analysed. Total numbers of parasites, number of parasites without beads, with beads bound, and with beads capped were quantified). Mean values of three independent experiments +/-SD were determined.

#### Live capping assays

Capping assay were adapted for live microscopy. Parasites were prepared as described above. After the addition of the diluted beads (5 µl of beads in 250 µl of H-H buffer) to the parasites, the dish was incubated for 10 minutes on ice. After incubation the media was exchanged for 500 µl of ice cold H-H buffer without beads. The dish was then directly transfer to the microscope. Time-lapse videos were taken with a 60X objective at 1 frame per second using a DeltaVision^®^ Core microscope. Analysis was made using ImageJ.

#### LPA, LPC, Bodipy and NGP uptake

LPA (Lyso-Phospatidic acid, TopFluor^®^ LysoPA, Avanti^®^ Polar Lipids, inc.), LPC (Lyso-Phospatidic choline, TopFluor^®^ LysoPC, Avanti^®^ Polar Lipids, inc.), bodipy (BODIPY™ 493/503, Thermo Fisher) and nanogold particles (NGP, Gold Nanoparticles 10 nM Cy5.5 labbeled, Nanocs) uptake assays were derived from the capping assays. Briefly, Ibidi live cell dishes (29 mm) were coated with 0.1% poly-L-Lysine for 30 min and washed with MilliQ water. NGP were diluted at 8 µl in 400 µl H-H buffer and left on ice for 30 min before use. LPA, LPC, and Bodipy were diluted to a concentration of 4µM in H-H buffer. Parasites were harvested, pelleted, and resuspended in cold H-H Buffer to achieve 10^7^ parasites/ml. Parasites were then transferred to poly-L-Lysine coated dishes and left on ice for 20 min. 250 µl of H-H buffer (± 4µM LPA, LPC or Bodipy) with or without 8 µl of diluted beads were added to the parasites. Immediately, the dish was incubated at 37 °C for 30 min. The experiment was stopped by addition of 2 ml of 4 % PFA and incubated at 4 °C for 10 min. The PFA was washed gently and parasite nuclei stained with Hoechst 0.01 %. For time course analysis, parasites were fixed at different time points (0, 1, 5, 10, 15, 20 and 30min) after the addition of the beads. For the drug and buffer assays, parasites were incubated for 10 min in the buffer of interest before their incubation on the coated dish. Mean values of three independent experiments +/-SD were determined.

#### SAG1 uptake

RH parasites were incubated with α-SAG1, 2µM LPA, and NGP on Ibidi live cell dishes (29 mm) coated with 0.1% poly-L-Lysine as described above. After 30 minutes incubation at 37°C, parasites were fixed and permeabilised. Secondary antibodies directed toward α-SAG1 were used to evaluate the uptake. Mean values of three independent experiments +/-SD were determined.

#### Secretion evaluation

RH Parasites were treated for 30 minutes with LPA (2 µM final) with or without particles as described above. After 30 min, the media was exchanged twice and parasite washed thoroughly to collect them and transfer them to a new dish with either H-H buffer (minimal media), Dulbecco’s modified Eagle’s medium (DMEM) supplemented with 10 % Foetal Bovine serum, 2 mM L-glutamine and 25 mg/ml gentamycin (complete media), or cover slips with host cell in complete media (Host cell). After 30 minutes of incubation, parasites were fixed and the presence of LPA and NGP were evaluated. For parasites on host cells, a dual SAG-1 (without permeabilization) and GAP45 (with permeabilization) was carried out to differentiate invaded from extracellular parasites. Mean values of three independent experiments +/-SD were determined and compared to a control fixed at the end of the first 30 minutes incubation with LPA and NGP (t=0).

#### Plaque assay

Parasite were treated for 30 minutes with or without LPA (2µM Final). 1×10^3^ parasites were inoculated on a confluent layer of HFFs and incubated for 5 days. After which, the HFFs were washed once with PBS and fixed with ice cold MeOH for 20 minutes. HFFs were stained with Giemsa with plaque area measured using Fiji software. Mean values of three independent experiments +/-SD were determined.

#### Immunofluorescence analysis

Carried out as previously described (8). Briefly, parasites were fixed in 4% paraformaldehyde for 10 min at 4°C. Afterwards, coverslips were blocked and permeabilised in 2% BSA & 0.2% Triton X–100 in PBS for 20 min. The staining was performed using the indicated combinations of primary antibodies for 1 h. Followed by the incubation with secondary AlexaFluor 350, AlexaFluor 488, AlexaFluor 594 or AlexaFluor 633 conjugated antibodies (1:3000, Invitrogen– Molecular Probes) for another 45 min, respectively. For quantification, mean values of three independent experiments +/-SD were determined.

#### SIM imaging

Super-Resolution Microscopy (SR-SIM) was carried out using an ELYRA PS.1 microscope (Zeiss). Images were acquired using a Plan Apochromat 63x, 1.4 NA oil immersion lens, recorded with a CoolSNAP HQ camera (Photometrics) and analysed using ZEN Black software (Zeiss) and ImageJ software.

#### Transmission electron microscopy

Extracellular parasites (±LPA/NGP see NGP uptake above) were fixed with 2.5% glutaraldehyde in 0.1 M phosphate buffer pH 7.4 after the indicated incubation. Samples were processed for routine electron microscopy as described previously (49) and examined in a JEOL 1200EX electron microscope.

#### Correlative light-electron microscopy (CLEM)

Uptake assays were carried out in gridded glass bottom petri dishes (MatTek). Parasites presenting clear LPA and NGP uptake were imaged with SR-SIM in an ELYRA PS.1 microscope (Carl Zeiss, Germany), and the material was fixed in 2.5% glutaraldehyde and 4% paraformaldehyde in 0.1 M phosphate buffer; and processed for transmission electron microscopy as described previously (49). Thin sections of the same areas imaged in 3D-SIM were imaged in a Tecnai T20 transmission electron microscope (FEI, Netherlands).

## Acknowledgements

We would like to thank Gurman Pall, Javier Perez, and Mario Del Rosario, as well as other members of the Meissner lab, for thoughtful discussions. We would also like to thank Prof. John Boothroyd (University of Stanford), Prof. Peter Bradley (University of California), Prof. Dominque Soldati-Favre (University of Geneva), Prof. Vern Carruthers (University of Michigan), and Dr Maryse Lebrun (University of Montpellier) for generously providing antibodies and parasite strains.

## Contributions

<u>Simon Gras:</u> Conceptualization, Data Curation, Formal Analysis, Investigation, Methodology, Project Administration, Writing – Original Draft Preparation, Writing – Review & Editing. <u>Elena Jimenez-Ruiz</u>: Conceptualization, Data Curation, Formal Analysis, Investigation, Methodology, Writing – Original Draft Preparation, Writing – Review & Editing. <u>Christen Klinger:</u> Formal Analysis, Investigation, Writing – Review & Editing. <u>Leandro Lemgruber</u>: Formal Analysis, Methodology. <u>Markus Meissner:</u> Conceptualization, Data Curation, Funding Acquisition, Writing – Original Draft Preparation, Writing – Review & Editing.

## Abbreviation

LPA: Lysophosphatidic acid
CDE: clathrin-dependant endocytosis
CDI: clathrin-independent endocytosis
CD: Cytochalasin-D
TFDC: Trifluoperazine di-hydrochloride
DN: Dominant negative
CHC: clathrin heavy chain
LPC: Lysophosphatidyl choline
BP: Bodipy
NGP: nanogold particles
VAC: vacuolar-like compartment
ER: endoplasmic reticulum
CLEM: corelative light and electron microscopy
TEM: transmission electron microscopy

## References

1. Lopes-Mori FM, Mitsuka-Bregano R, Capobiango JD, Inoue IT, Reiche EM, Morimoto HK, et al. Programs for control of congenital toxoplasmosis. Rev Assoc Med Bras (1992). 2011;57(5):594–9.

2. Kafsack BF, Beckers C, Carruthers VB. Synchronous invasion of host cells by Toxoplasma gondii. Mol Biochem Parasitol. 2004;136(2):309–11.

3. Meissner M, Ferguson DJ, Frischknecht F. Invasion factors of apicomplexan parasites: essential or redundant? Curr Opin Microbiol. 2013;16(4):438–44.

4. Bretscher MS. Asymmetry of single cells and where that leads. Annu Rev Biochem. 2014;83:275–89.

5. Hegge S, Munter S, Steinbuchel M, Heiss K, Engel U, Matuschewski K, et al. Multistep adhesion of Plasmodium sporozoites. Faseb J. 2010;24(7):2222–34.

6. Munter S, Sabass B, Selhuber-Unkel C, Kudryashev M, Hegge S, Engel U, et al. Plasmodium sporozoite motility is modulated by the turnover of discrete adhesion sites. Cell Host Microbe. 2009;6(6):551–62.

7. Whitelaw JA, Latorre-Barragan F, Gras S, Pall GS, Leung JM, Heaslip A, et al. Surface attachment, promoted by the actomyosin system of Toxoplasma gondii is important for efficient gliding motility and invasion. BMC Biol. 2017;15(1):1.

8. Egarter S, Andenmatten N, Jackson AJ, Whitelaw JA, Pall G, Black JA, et al. The toxoplasma Acto-MyoA motor complex is important but not essential for gliding motility and host cell invasion. PLoS One. 2014;9(3):e91819.

9. Maritzen T, Schachtner H, Legler DF. On the move: endocytic trafficking in cell migration. Cell Mol Life Sci. 2015;72(11):2119–34.

10. Tanaka M, Kikuchi T, Uno H, Okita K, Kitanishi-Yumura T, Yumura S. Turnover and flow of the cell membrane for cell migration. Sci Rep. 2017;7(1):12970.

11. Quadt KA, Streichfuss M, Moreau CA, Spatz JP, Frischknecht F. Coupling of Retrograde Flow to Force Production During Malaria Parasite Migration. ACS Nano. 2016;10(2):2091–102.

12. Carruthers V, Boothroyd JC. Pulling together: an integrated model of Toxoplasma cell invasion. Curr Opin Microbiol. 2007;10(1):83–9.

13. Jimenez-Ruiz E, Morlon-Guyot J, Daher W, Meissner M. Vacuolar protein sorting mechanisms in apicomplexan parasites. Mol Biochem Parasitol. 2016.

14. Hakansson S, Morisaki H, Heuser J, Sibley LD. Time-lapse video microscopy of gliding motility in Toxoplasma gondii reveals a novel, biphasic mechanism of cell locomotion. Mol Biol Cell. 1999;10(11):3539–47.

15. McGovern OL, Rivera-Cuevas Y, Kannan G, Narwold AJ, Jr., Carruthers VB. Intersection of endocytic and exocytic systems in Toxoplasma gondii. Traffic. 2018;19(5):336–53.

16. Dou Z, McGovern OL, Di Cristina M, Carruthers VB. Toxoplasma gondii ingests and digests host cytosolic proteins. MBio. 2014;5(4):e01188–14.

17. Kawauchi T. Cell adhesion and its endocytic regulation in cell migration during neural development and cancer metastasis. Int J Mol Sci. 2012;13(4):4564–90.

18. Gao W, Shi P, Chen X, Zhang L, Liu J, Fan X, et al. Clathrin-mediated integrin alphaIIbbeta3 trafficking controls platelet spreading. Platelets. 2018;29(6):610–21.

19. Doherty GJ, McMahon HT. Mechanisms of endocytosis. Annu Rev Biochem. 2009;78:857–902.

20. Woo YH, Ansari H, Otto TD, Klinger CM, Kolisko M, Michalek J, et al. Chromerid genomes reveal the evolutionary path from photosynthetic algae to obligate intracellular parasites. Elife. 2015;4.

21. Pieperhoff MS, Schmitt M, Ferguson DJ, Meissner M. The role of clathrin in post-Golgi trafficking in Toxoplasma gondii. PLoS One. 2013;8(10):e77620.

22. Kremer K, Kamin D, Rittweger E, Wilkes J, Flammer H, Mahler S, et al. An overexpression screen of Toxoplasma gondii Rab-GTPases reveals distinct transport routes to the micronemes. PLoS Pathog. 2013;9(3):e1003213.

23. Breinich MS, Ferguson DJ, Foth BJ, van Dooren GG, Lebrun M, Quon DV, et al. A dynamin is required for the biogenesis of secretory organelles in Toxoplasma gondii. Curr Biol. 2009;19(4):277–86.

24. Agop-Nersesian C, Egarter S, Langsley G, Foth BJ, Ferguson DJ, Meissner M. Biogenesis of the Inner Membrane Complex Is Dependent on Vesicular Transport by the Alveolate Specific GTPase Rab11B. PLoS Pathog. 2010;6(7):e1001029.

25. Agop-Nersesian C, Naissant B, Ben Rached F, Rauch M, Kretzschmar A, Thiberge S, et al. Rab11A-controlled assembly of the inner membrane complex is required for completion of apicomplexan cytokinesis. PLoS Pathog. 2009;5(1):e1000270.

26. Miranda K, Pace DA, Cintron R, Rodrigues JC, Fang J, Smith A, et al. Characterization of a novel organelle in Toxoplasma gondii with similar composition and function to the plant vacuole. Mol Microbiol. 2010;76(6):1358–75.

27. Parussini F, Coppens I, Shah PP, Diamond SL, Carruthers VB. Cathepsin L occupies a vacuolar compartment and is a protein maturase within the endo/exocytic system of Toxoplasma gondii. Mol Microbiol. 2010;76(6):1340–57.

28. King CA. Cell surface interaction of the protozoan Gregarina with concanavalin A beads-implications for models of gregarine gliding. Cell Biol Int Rep. 1981;5(3):297–305.

29. Frenal K, Dubremetz JF, Lebrun M, Soldati-Favre D. Gliding motility powers invasion and egress in Apicomplexa. Nature reviews Microbiology. 2017;15(11):645–60.

30. Dutta D, Donaldson JG. Search for inhibitors of endocytosis: Intended specificity and unintended consequences. Cell Logist. 2012;2(4):203–8.

31. Daniel JA, Chau N, Abdel-Hamid MK, Hu L, von Kleist L, Whiting A, et al. Phenothiazine-derived antipsychotic drugs inhibit dynamin and clathrin-mediated endocytosis. Traffic. 2015;16(6):635–54.

32. Mickler FM, Mockl L, Ruthardt N, Ogris M, Wagner E, Brauchle C. Tuning nanoparticle uptake: live-cell imaging reveals two distinct endocytosis mechanisms mediated by natural and artificial EGFR targeting ligand. Nano Lett. 2012;12(7):3417–23.

33. Moolenaar WH. Lysophosphatidic acid signalling. Curr Opin Cell Biol. 1995;7(2):203–10.

34. Kuriyama S, Theveneau E, Benedetto A, Parsons M, Tanaka M, Charras G, et al. In vivo collective cell migration requires an LPAR2-dependent increase in tissue fluidity. J Cell Biol. 2014;206(1):113–27.

35. Urs NM, Jones KT, Salo PD, Severin JE, Trejo J, Radhakrishna H. A requirement for membrane cholesterol in the beta-arrestin-and clathrin-dependent endocytosis of LPA1 lysophosphatidic acid receptors. J Cell Sci. 2005;118(Pt 22):5291–304.

36. Harper JM, Hoff EF, Carruthers VB. Multimerization of the Toxoplasma gondii MIC2 integrin-like A-domain is required for binding to heparin and human cells. Mol Biochem Parasitol. 2004;134(2):201–12.

37. Jacot D, Tosetti N, Pires I, Stock J, Graindorge A, Hung YF, et al. An Apicomplexan Actin-Binding Protein Serves as a Connector and Lipid Sensor to Coordinate Motility and Invasion. Cell Host Microbe. 2016;20(6):731–43.

38. Calderwood DA, Ginsberg MH. Talin forges the links between integrins and actin. Nat Cell Biol. 2003;5(8):694–7.

39. Barnhart E, Lee KC, Allen GM, Theriot JA, Mogilner A. Balance between cell-substrate adhesion and myosin contraction determines the frequency of motility initiation in fish keratocytes. Proc Natl Acad Sci U S A. 2015;112(16):5045–50.

40. Fogelson B, Mogilner A. Computational estimates of membrane flow and tension gradient in motile cells. PLoS One. 2014;9(1):e84524.

41. Keren K, Yam PT, Kinkhabwala A, Mogilner A, Theriot JA. Intracellular fluid flow in rapidly moving cells. Nat Cell Biol. 2009;11(10):1219–24.

42. Stroka KM, Jiang H, Chen SH, Tong Z, Wirtz D, Sun SX, et al. Water permeation drives tumor cell migration in confined microenvironments. Cell. 2014;157(3):611–23.

43. Lammermann T, Bader BL, Monkley SJ, Worbs T, Wedlich-Soldner R, Hirsch K, et al. Rapid leukocyte migration by integrin-independent flowing and squeezing. Nature. 2008;453(7191):51–5.

44. Russell DG, Sinden RE. The role of the cytoskeleton in the motility of coccidian sporozoites. J Cell Sci. 1981;50:345–59.

45. Andenmatten N, Egarter S, Jackson AJ, Jullien N, Herman JP, Meissner M. Conditional genome engineering in Toxoplasma gondii uncovers alternative invasion mechanisms. Nat Methods. 2013;10(2):125–7.

46. Rugarabamu G, Marq JB, Guerin A, Lebrun M, Soldati-Favre D. Distinct contribution of Toxoplasma gondii rhomboid proteases 4 and 5 to micronemal protein protease 1 activity during invasion. Mol Microbiol. 2015;97(2):244–62.

47. Shen B, Buguliskis JS, Lee TD, Sibley LD. Functional analysis of rhomboid proteases during Toxoplasma invasion. MBio. 2014;5(5):e01795–14.

48. Gras S, Jackson A, Woods S, Pall G, Whitelaw J, Leung JM, et al. Parasites lacking the micronemal protein MIC2 are deficient in surface attachment and host cell egress, but remain virulent in vivo. Wellcome Open Res. 2017;2:32.

49. Periz J, Whitelaw J, Harding C, Gras S, Del Rosario Minina MI, Latorre-Barragan F, et al. Toxoplasma gondii F-actin forms an extensive filamentous network required for material exchange and parasite maturation. Elife. 2017;6.

50. Wilson BJ, Allen JL, Caswell PT. Vesicle trafficking pathways that direct cell migration in 3D and in vivo. Traffic. 2018.

51. Keren K. Cell motility: the integrating role of the plasma membrane. Eur Biophys J. 2011;40(9):1013–27.

52. Kay RR, Langridge P, Traynor D, Hoeller O. Surface area regulation: underexplored yet crucial in cell motility. Nature Reviews Molecular Cell Biology. 2008;9:662.

53. Bretscher MS. Getting membrane flow and the cytoskeleton to cooperate in moving cells. Cell. 1996;87(4):601–6.

54. Stadler RV, White LA, Hu K, Helmke BP, Guilford WH. Direct measurement of cortical force generation and polarization in a living parasite. Mol Biol Cell. 2017;28(14):1912–23.

55. Tomavo S. Evolutionary repurposing of endosomal systems for apical organelle biogenesis in Toxoplasma gondii. Int J Parasitol. 2014;44(2):133–8.

56. Gaits F, Fourcade O, Le Balle F, Gueguen G, Gaige B, Gassama-Diagne A, et al. Lysophosphatidic acid as a phospholipid mediator: pathways of synthesis. FEBS Lett. 1997;410(1):54–8.

57. Antonescu CN, Danuser G, Schmid SL. Phosphatidic acid plays a regulatory role in clathrin-mediated endocytosis. Mol Biol Cell. 2010;21(16):2944–52.

58. Norambuena A, Metz C, Jung JE, Silva A, Otero C, Cancino J, et al. Phosphatidic acid induces ligand-independent epidermal growth factor receptor endocytic traffic through PDE4 activation. Mol Biol Cell. 2010;21(16):2916–29.

59. Roth MG. Molecular mechanisms of PLD function in membrane traffic. Traffic. 2008;9(8):1233–9.

60. Bullen HE, Jia Y, Yamaryo-Botte Y, Bisio H, Zhang O, Jemelin NK, et al. Phosphatidic Acid-Mediated Signaling Regulates Microneme Secretion in Toxoplasma. Cell Host Microbe. 2016;19(3):349–60.

61. Nichols BA, Chiappino ML, Pavesio CE. Endocytosis at the micropore of Toxoplasma gondii. Parasitol Res. 1994;80(2):91–8.

62. Brown KM, Long S, Sibley LD. Conditional Knockdown of Proteins Using Auxin-inducible Degron (AID) Fusions in Toxoplasma gondii. Bio Protoc. 2018;8(4).

63. Tomavo S, Slomianny C, Meissner M, Carruthers VB. Protein trafficking through the endosomal system prepares intracellular parasites for a home invasion. PLoS Pathog. 2013;9(10):e1003629.

64. Sidik SM, Huet D, Ganesan SM, Huynh MH, Wang T, Nasamu AS, et al. A Genome-wide CRISPR Screen in Toxoplasma Identifies Essential Apicomplexan Genes. Cell. 2016;166(6):1423–35 e12.

65. Donald RG, Carter D, Ullman B, Roos DS. Insertional tagging, cloning, and expression of the Toxoplasma gondii hypoxanthine-xanthine-guanine phosphoribosyltransferase gene. Use as a selectable marker for stable transformation. J Biol Chem. 1996;271(24):14010–9.

66. Huynh MH, Carruthers VB. Tagging of endogenous genes in a Toxoplasma gondii strain lacking Ku80. Eukaryot Cell. 2009;8(4):530–9.

67. Sloves PJ, Delhaye S, Mouveaux T, Werkmeister E, Slomianny C, Hovasse A, et al. Toxoplasma Sortilin-like Receptor Regulates Protein Transport and Is Essential for Apical Secretory Organelle Biogenesis and Host Infection. Cell Host Microbe. 2012;11(5):515–27.

